# Genotype-specific developmental plasticity shapes the timing and robustness of reproductive capacity in *Caenorhabditis elegans*

**DOI:** 10.1101/045609

**Authors:** Bradly Alicea

**Affiliations:** University of Illinois, Urbana-Champaign.

**Keywords:** Developmental Plasticity, Computational Biology, Transgenerational Plasticity, Evolution of Development, Reproductive Dynamics

## Abstract

The effects of environmental stress on developmental phenotypes allows us to observe the adaptive capacity of various genotypes. In single-genotype populations, this allows for observing the effects of standing variation on the process of developmental plasticity. Typically, such efforts do not allow us to distinguish the effect of genotype from the effects of environment. In the Nematode *Caenorhabditis elegans*, however, the L1 stage of development allows for precise and controlled imposition of starvation on a synchronized population of individuals. This allows for systematic comparisons within and between defined genotypes.

This study provides a number of innovations in the study of Nematode developmental plasticity. The first is to demonstrate systems-level mechanisms of adaptive response for various genetic mutant phenotypes to extended L1 arrest. The effects of developmental deprivation are quantifiable by applying statistical modeling techniques to reproductive time-series. This is characterized for both a wildtype strain (N2) and a host of genetic mutant strains. A further contribution is investigating the effects of starvation at multi-generational timescales. This includes both two generations post-starvation and multiple starvation experiences with selection for fecundity over 11 generations.

Statistical techniques such as Kernel Density Estimation (KDE), fold-change analysis, and Area Under the Curve (AUC) analysis reveal differences from baseline in the form of shifts in reproductive timing and kurtosis in peak reproductive capacity characterize adaptive responses that are semi-independent of environment. Extending these patterns of response to *ad hoc* comparisons with multi-generational contexts reveals that different genotypes characterized by mutations of specific genetic loci lead to vastly different outcomes.

## INTRODUCTION

Developmental plasticity has been defined as a universal property of phenotypes that reveal intra-individual variation due to both environmental and genetic influences (West-Eberhard, 2003). Plasticity also allows us to explore the adaptability of a specific genotype, or to what degree these genotypes express novel phenotypes in response to environmental selection (Moczek et.al,2011). The Nematode *Caenorhabditis elegans* provides us with a window into developmental plasticity that allows us to easily partition the genetic and environmental components of this process.

There are two periods of diapause in *C. elegans* developmental life-history which provide opportunities to observe divergent developmental trajectories resulting from environmental conditions: extended L1 arrest (Johnson et.al, 1984) and dauer arrest (Burnell et.al, 2005). While the dauer stage results in a set of immediate facultative phenotypic adaptations that prolong the lifespan (Wolkow et.al, 2000), extended arrest during the L1 stage results in facultative adaptations that persist throughout subsequent developmental stages after recovery (Kang and Avery, 2009; Artyukhin and Avery, 2013). Often characterized as a stress response to starvation, the adaptations related to recovery from L1 arrest is not immediate and does not result in an alternative phenotype. However, it does act to influence the development of individual organisms in a number of fundamental ways. According to Lee et.al (2012), extended L1 arrest represents starvation conditions that have a metabolic cost that adversely affects an individual’s subsequent development and reproductive capacity. The Lee et.al (2012) study introduces the notion of L1 longevity, which requires the organism to maintain a low level of metabolism during the course of L1 arrest. While this can have immediate benefits to an organism’s survival, this should disproportionally affect mutant strains such as AMPK and *daf-c*, which have defects for metabolic regulation and insulin signaling that could affect their reproductive response to extended L1 arrest (Apfeld et.al, 2004; Michaelson et.al, 2010). Furthermore, extended L1 arrest can lead to damaged gonads (Lee et.al, 2012) in general and reduced viability and failure to maintain mitotic quiescence in germline stem cells in AMPK mutants specifically (Fukuyama et.al, 2012). Such dysregulation of the developmental process can further affect the fecundity of starvation-experiencing individuals.

The ability to produce synchronized populations with starvation experience during extended L1 arrest also provides a window into the timing of development and reproduction, which in turn allows us to assess the effects of characterized mutations on plasticity exhibited by specific individuals in a population. Baugh (2013) suggests that extended L1 arrest can lead to changes in the heterochronic trajectory of individual worms representing individual phenotypes. One estimation suggests that 24h period of extended L1 arrest results in a 3h delay in normal developmental progression (Lee et.al, 2012). Both of these effects are contingent upon genotypic background, which may mitigate or exaggerate these responses. Aside from specific functional effects expected amongst AMPK and *daf-c* mutants, certain *daf-d* mutants can also demonstrate effects related to developmental rate and reproduction. As the regulation of developmental rate is a daf-16/FOXO-dependent process (Murphy and Hu, 2013), daf-16 mutant genotypes should also be affected by extended L1 arrest.

Starvation during extended L1 arrest also triggers adaptive responses at the molecular level. In Maxwell et.al (2012), it is shown using RNA-seq that a variety of mRNAs representing diverse regulatory functions are upregulated during a window of one to six hours after recovery from extended L1 arrest. Gene expression during further characterized in terms of the availability of transcriptional resources during recovery by a modeling study (Zaslaver, Baugh, and Sternberg, 2011), in which moderate increases in transcriptional resources are facilitated by temporarily increased rates of transcription amongst a diverse set of genes and regulatory elements. In turn, this set of molecular processes allow for rapid growth and recovery of the phenotype.

Finally, there are a number of transgenerational effects resulting from starvation during extended L1 arrest. These include reduced fecundity, slower growth, and fewer offspring. In general, previously starved larvae are more sensitive to subsequent starvation, although individuals most severely affected by extended L1 arrest also tend to produce more robust offspring (Jobson et.al, 2015). Rechavi et.al (2014) demonstrate that F3 descendents of worms experiencing extended L1 arrest show an increased lifespan, inheritance of small RNAs, and other adaptive traits. While it is unclear whether or not lifespan is linked to a shift in reproduction to later in life-history is an evolutionary trade-off (2007), there may be an effect of "decanalization" (Jobson et.al, 2015), in which some individuals develop and reproduce more quickly, while other individuals develop and reproduce more slowly.

### Initial Hypotheses

This paper will address three hypotheses, each of which address different components of developmental plasticity. These hypotheses address various aspects of reproductive capacity, defined as the developmental precursors and physiological functionality of producing viable offspring.

**H1:** the experience of starvation during extended L1 arrest leads to developmental plasticity expressed as mutant genotype-dependent changes in lifetime fecundity.
**H2:** the experience of starvation during extended L1 arrest leads to developmental plasticity expressed as delays, accelerations, intensifications, and dampening of reproductive capacity in a genetic pathway-dependent manner.
**H3:** the experience of starvation during extended L1 arrest has multi-generational effects at both short time-scales (two generations) and longer time-scales. At longer time-scales, the compensatory responses of developmental plasticity are expressed in genealogically dependent manner.

By combining experimental techniques with mathematical modeling, we can gain insight into reproductive capacity and developmental plasticity associated with extended L1 arrest across a series of *C. elegans* mutant genetic strains. This generalized effect is unique within and between several different classes of mutant over five days of reproductive history. Using resampled representations of fecundity measurements, we can characterize the timing, magnitude, and overall shape of reproductive response. This may also have implications for evolutionary selection, as the data for starvation resulting from extended L1 arrest is compared to similar results over multiple generations. This includes both short-term heritable effects and longer-term effects on heritability and demography.

## RESULTS

This analysis will be organized around several ways of testing our three initial hypotheses. First, we would like to know to what degree developmental plasticity leads to changes in reproductive capacity and timing. To do this, the hanging drop method (see Methods) is used to both observe per day reproduction and limit potential maternal effects of all eggs laid to a single hermaphroditic ancestor (Harvey and Orbidans, 2011). We would then like to know whether or not the magnitude of these changes in reproductive capacity and timing are contingent upon genetic background, particularly as they relate to known functional mutations. This is done by interpolating the data into reproductive functions (see Methods). Reproductive functions are utilized to investigate reproductive capacity and developmental plasticity in AMPK, *daf*, and AMPK/*daf-c* mutants, the last of which provides a hybrid genetic background. After exploring the potential for developmental plasticity in AMPK and AMPK/*daf-c* mutants, we then conduct a more extensive investigation of *daf-c*, *daf-d*, and *daf-c*/*daf-d* mutant genotypes. The differences in reproductive functions between extended L1 arrest-experiencing individuals and non-experienced individuals is assessed quantitatively using an approach called developmental delay analysis. Finally, we explore the multigenerational effects of extended L1 arrest in AMPK and AMPK/*daf-c* mutant strains at two timescales (2 and 11 generations).

### Histograms of Net Fecundity by Genotype Group

Initially, we sought to establish a connection between L1 arrest, developmental delay, and a mechanism for changes in reproductive fitness. To better understand the diversity of responses across life-history for specific classes of genotype, we visualized the fecundity of multiple groups of genetic strain using kernel-density estimation (KDE). KDE provides us with a non-parametric description of distributions within and between populations that serve as an approximation of reproductive fitness. Using the reproductive capacity across the adult lifespan of single worms representing several classes of defined mutation, the KDE analyses is used to summarize net fecundity (see Methods). We contrast each set of non-L1 arrested individuals with each set of L1-arrested individuals. The reproductive fitness distribution for several classes of genotype is shown in Figure S1. Each histogram in Figure S1 shows the distribution of reproductive capacity for a population when the effects of developmental arrest are subtracted from the reproductive capacity of individuals experiencing no developmental arrest.

### Approximated Time-course Analysis

To better understand these results in the context of reproductive life-history, we measured and mathematically approximated the reproductive time-course (5d) for both L1 arrest-experiencing and non-L1 arrest-experiencing individuals (Figure 1). For the wild-type genotype (N2), developmental arrest does lead to two developmental-related effects. One effect of developmental arrest is a dampening of peak reproductive fecundity, which is 1.61-fold less than during normal development. A dampening in fecundity suggests that L1 arrest affects vulval development in a specific genetic background. The second effect is the relative onset of egg-laying leading to viable offspring, which in the case of Figure 1 (N2) is delayed by 1-2 days. A temporal delay in peak fecundity suggests that L1 arrest affects the developmental timing of sperm production and/or a change in egg-laying.

**Figure 1.**
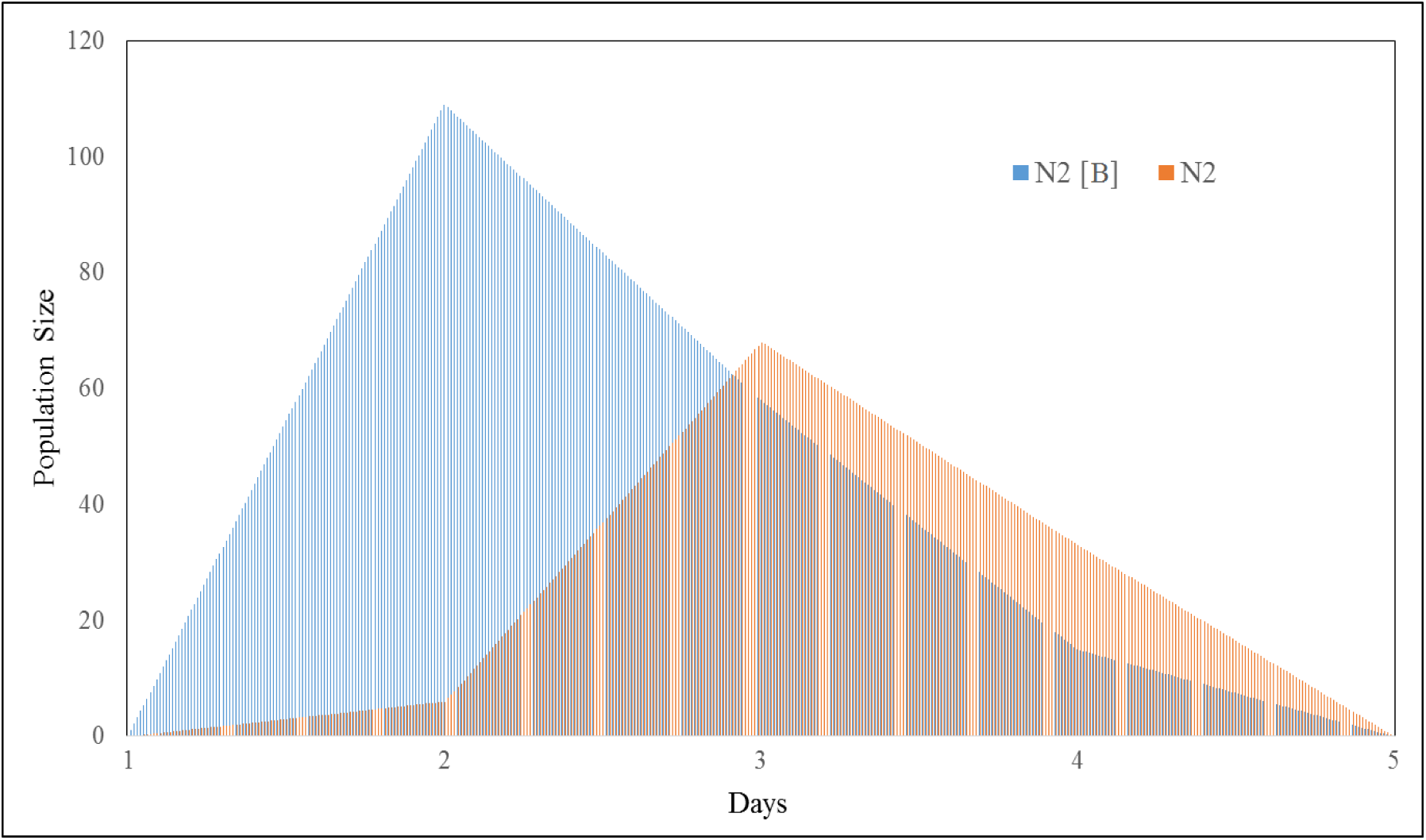
Graphical comparison of the wild-type genotype (N2) for two environmental conditions: normal development (blue) versus developmental arrest (orange/pink).

**AMPK mutants.** Fold-change differences are also calculated to more directly compare the time and maximum fecundity of each genotype. This is done by taking the first moment of the distributions shown in Figure 1. The unimodal peak represents the peak fecundity, which can be contrasted with various arrest treatments as well as compared across genotypes. Differences in fecundity are characterized by the relative difference in height between different conditions. The difference in time between these peaks can also be calculated, yielding a relative measure of developmental delay. Table 1 shows these calculations for a wild-type genotype and three AMPK mutants (aak-1, aak-2, and aak-1; aak-2).

**Table 1.** Fold-change differences between baseline and extended L1 arrest conditions for wildtype (N2) and three different AMPK mutant genotypes. Fold-change difference was calculated on the peak fecundity across replicates and its time-point of occurrence (within 5d of L4).

| Genotype * Environment | Peak Fecundity | Peak Time (days) | Fecundity fold-difference | Time fold-difference |
| --- | --- | --- | --- | --- |
| N2 | 68 | 3.0 | -1.61 | -1.5 |
| N2 (control) | 109 | 2.0 |  |  |
| *aak-1* | 29 | 2.0 | -2.31 | 0 |
| *aak-1* (control) | 67 | 2.0 |  |  |
| aak-2 | 159 | 5.0 | 1.63 | -2.1 |
| aak-2 (control) | 98 | 2.42 |  |  |
| aak-1; aak-2 | 48 | 2.0 | -1.21 | 1.225 |
| aak-1; aak-2 (control) | 58 | 2.45 |  |  |

Now that we have established that experience of L1 arrest can lead to arrest in reproductive development and a loss of fecundity, we can now examine the effects of L1 arrest on reproductive development and fecundity in a mutant genotype. For our first demonstration, we have chosen mutations in the AMPK pathway. Figure 2 (A-C) shows a graphical comparison of normal development versus developmental arrest for our AMPK mutants in a manner identical to Figure 1. Figure 2 (D) shows a comparison of all three AMPK mutants and their performance after developmental arrest.

**Figure 2.**
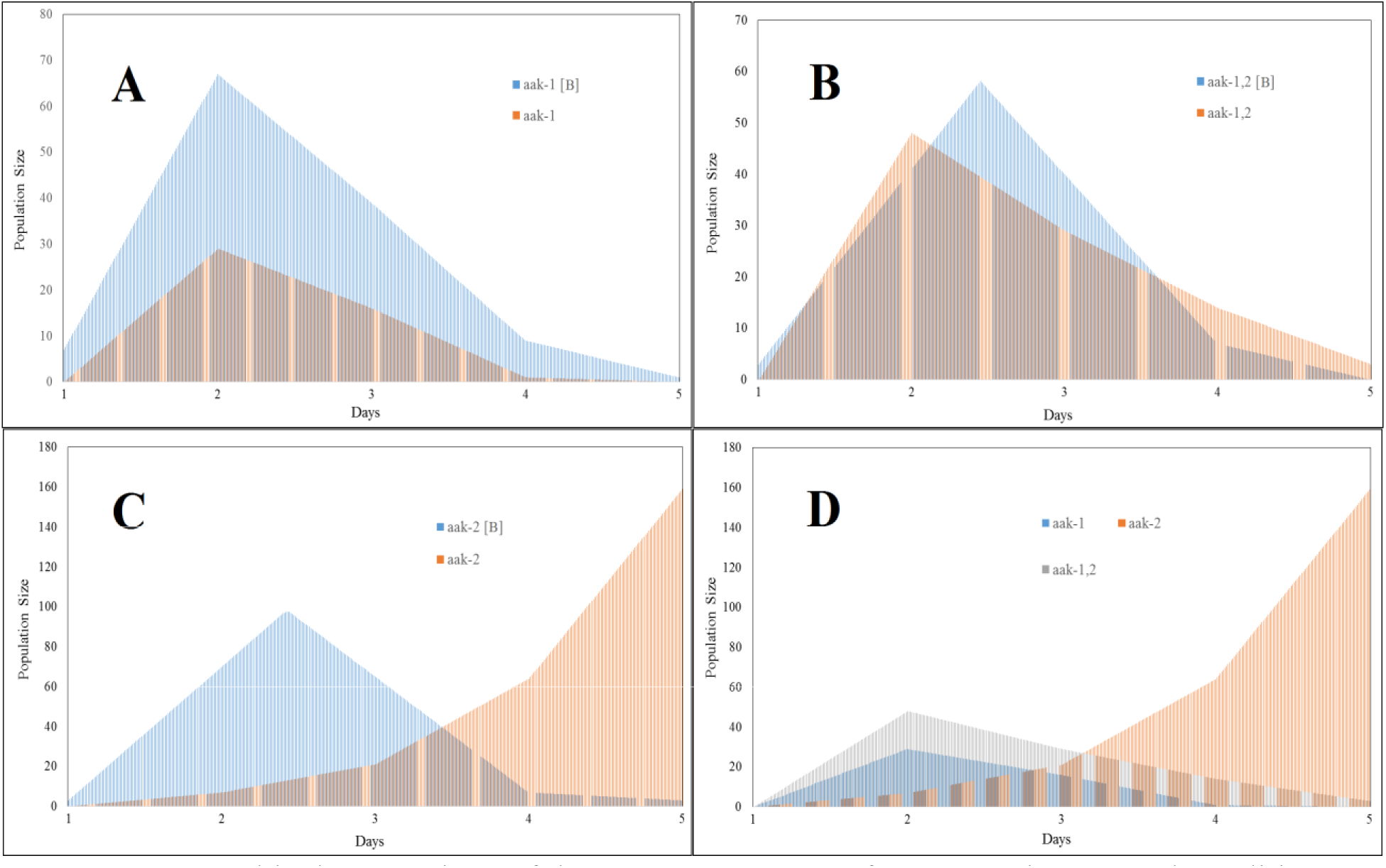
Graphical comparison of the AMPK genotypes for two environmental conditions: normal development (blue) versus developmental arrest (orange/pink). A: comparison for aak-1, B: comparison for aak-1; aak-2, C: comparison for aak-2, D: comparison of developmental arrest effects for all three AMPK mutant strains.

For the double mutant (*aak-1*; *aak-2*, alleles *tm*1944 and *ok*524, respectively), there is both a slight decrease in peak fecundity with respect to baseline and a more significant decrease with respect to baseline for the *aak-1* (allele tm1944) single mutant. There does not appear to be a developmental lag in neither the *aak-1* mutant nor the double mutant (*aak-1*; *aak-2*). The increase of peak reproductive fecundity in the double mutant is 1.23-fold greater than during normal development (Table 1), while there is neither delay nor acceleration with respect to the timing of peak fecundity.

Of particular note is the effect of the *aak-2* mutation (allele ok524), which appears to have a compensatory effect for experiencing extended L1 arrest. We can say this by looking at both an increase in peak fecundity between baseline and the treatment conditions (1.63-fold) in addition to a very long developmental lag (-2.1-fold) in the treatment condition. Moreover, there is a large increase in offspring production late in the lifespan based on incremental increases in fecundity up to 5d. The shape of the function for *aak-2* individuals that have experienced extended L1 arrest suggests a very long developmental delay, which could be distinct from a simple dampened and delayed response (e.g. isometric with respect to baseline).

#### AMPK/*daf-c* mutants

We can also look at the pairwise differences in both developmental (Table S2) and demographic lag (Table S3) to understand more subtle differences between various mutant strains. The bivariate relationship between these two parameters may yield other interesting trends (Figure S4). While deficits in metabolic regulation are known for AMPK mutants, the developmental effects are not well-characterized. Genetic crosses inspired by Fukuyama et.al (2012) between an AMPK double-mutant (*aak-1*; *aak-2*) and a **daf-c** mutant (*daf-7*) may provide better information regarding interactions between metabolic regulation and the mechanisms of larval development,

As the AMPK mutants exhibit lower reproductive robustness overall, we can also examine this mutant genetic background can also be combined with the *daf-c* mutant background (Table 2). Given their direct effects on dauer formation, *daf-c* mutants might provide us with a more direct analogue between developmental perturbation (L1 arrest) and developmental robustness (adaptive plasticity). We will first examine the fecundity and developmental trajectory of a hybrid genetic background (AMPK/daf-c mutants), and then examine these features in a variety of *daf-c*, *daf-d* and *daf-c*/*daf-d* mutants.

**Table 2.** Fold-change differences between baseline and extended L1 arrest conditions for a representative *daf-c* strain (daf-7) and three different AMPK/*daf-c* mutant genotypes. Fold-change difference was calculated on the peak fecundity across replicates and its time-point of occurrence (within 5d of L4).

| Genotype *<br>Environment | Peak<br>Fecundity | Peak Time<br>(days) | Fecundity fold-difference | Time fold-difference |
| --- | --- | --- | --- | --- |
| *daf-7* | 42.32 | 3.6 | -6.59 | -1.06 |
| *daf-7* (control) | 278.8 | 3.4 |  |  |
| *aak-1; daf-7* | 91 | 4.0 | -4.11 | 0 |
| *aak-1; daf-7*<br>(control) | 374 | 4.0 |  |  |
| *aak-2; daf-7* | 180 | 4.0 | -1.82 | 0 |
| *aak-2; daf-7*<br>(control) | 328 | 4.0 |  |  |
| *aak-1; aak-2; daf-7* | 220 | 4.0 | -1.42 | 0 |
| *aak-1; aak-2; daf-7*<br>(control) | 312 | 4.0 |  |  |

Figure 3(A-C) shows a comparison between the three different AMPK/daf-c mutants in a manner identical to Figure 2. Figure 3 (D) shows a comparison between all three treatment conditions. In all three genotypes, there is both a delay in peak fecundity and a loss of fecundity (Table 2) after experiencing extended L1 arrest. In the case of Figure 3B and 3C (*aak-1*; *aak2*; *daf-7* and *aak-1*; *daf-7*), there is a deformation of the function for the treatment condition consistent with the baseline measurement. As in the case of the wildtype (N2), this translates to roughly a 1d delay in the normal developmental trajectory. In aak-1; daf-7, we show a steady increase in fecundity to peak fecundity with some delay relative to baseline. for aak-1; aak-2; daf-7, we see a more pronounced delay with a peak fecundity closer to that of the baseline. An even more pronounced delay is shown for *aak-2*; *daf-7*, with a sharper rise in peak fecundity. It is of note that the compensatory effects of the *aak-2* mutation alone are not replicated in the *aak-2*; *daf-7* mutants.

**Figure 3.**
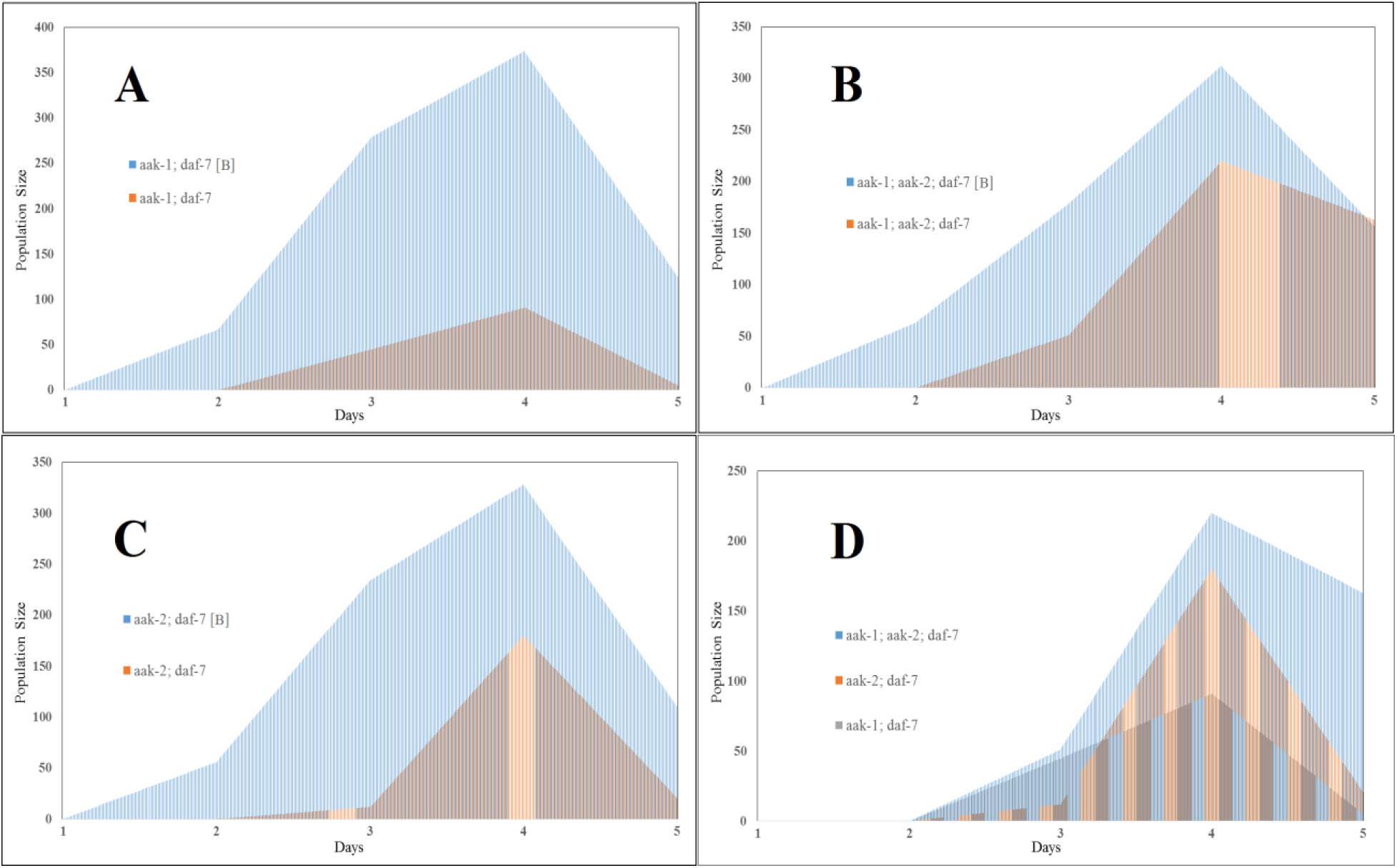
Graphical comparison of the AMPK/*daf-c* genotypes for two environmental conditions: normal development (blue) versus developmental arrest (orange/pink). A: comparison for *aak-1*; *daf-7*, B: comparison for *aak-1*; *aak-2*; *daf-7*, C: comparison for aak-2; daf-7, D: comparison of developmental arrest effects for all three AMPK/*daf-c* mutant strains.

#### *daf-c* and *daf-d* mutants

Moving on to *daf-c* and *daf-d* mutant genetic backgrounds, we can ask if mutations in genes in the dauer-formation pathway effect the reproductive dynamics of organisms that have experienced developmental arrest. This set of analyses excludes mutations in the AMPK pathway altogether, and addresses the effects of starvation during extended L1 arrest on mutations in the dauer-formation (*daf-c*) and dauer defective (*daf-d*) pathways. Shown in Table 3 are the differences between peak fecundity and peak time for the baseline and extended L1 arrest. Figure 4 shows the difference between extended L1 arrest and baseline for two *daf-c* mutants (*daf-7* and *daf-9*) and a single *daf-c*/*daf-d* mutant (*daf-7*; *daf-16(mu86)*). Two daf-c/daf-d mutants were used in Figures 4 and 5 to represent a partial dauer state produced through epistatic interactions first proposed by Vowels and Thomas (1992). In Figure 4A and B (*daf-c* mutants), there is a large difference in the kurtosis of peak fecundity between extended L1 arrest and baseline. In Figure 4C, there is a shift in peak fecundity to later in life-history, but no change in the kurtosis of peak fecundity.

**Table 3.** Fold-change differences between baseline and extended L1 arrest conditions for a wildtype strain (N2), two representative daf-d strains (daf-7 and daf-9), and two daf-c/daf-d strains (daf-7; daf-16(m27) and daf-7; daf-16(mu86)). Fold-change difference was calculated on the peak fecundity across replicates and its time-point of occurrence (within 5d of L4).

| Genotype *<br>Environment | Peak<br>Fecundity | Peak Time<br>(days) | Fecundity fold-difference | Time fold-difference |
| --- | --- | --- | --- | --- |
| N2 | 68 | 3.0 | -1.61 | -1.5 |
| N2 (control) | 109 | 2.0 |  |  |
| *daf-7* | 42.32 | 3.6 | -6.59 | -1.06 |
| *daf-7* (control) | 278.8 | 3.4 |  |  |
| *daf-9* | 11 | 4.0 | -8.72 | 0 |
| *daf-9* (control) | 76 | 4.0 |  |  |
| *daf-7; daf-16 (mu86)* | 114 | 4.0 | -1.24 | -1.19 |
| *daf-7; daf-16 (mu86)*<br>(control) | 141.4 | 3.35 |  |  |
| *daf-7; daf-16 (m27)*<br>(control)* | 112 | 4.0 | ----- | ----- |
| *daf-16 (mu86)*<br>(control)* | 297.04 | 2.39 | ----- | ----- |
| *daf-16 (m27)*<br>(control)* | 359 | 2 | ----- | ----- |

\* developmental arrest yielded no viable individuals.

**Figure 4.**
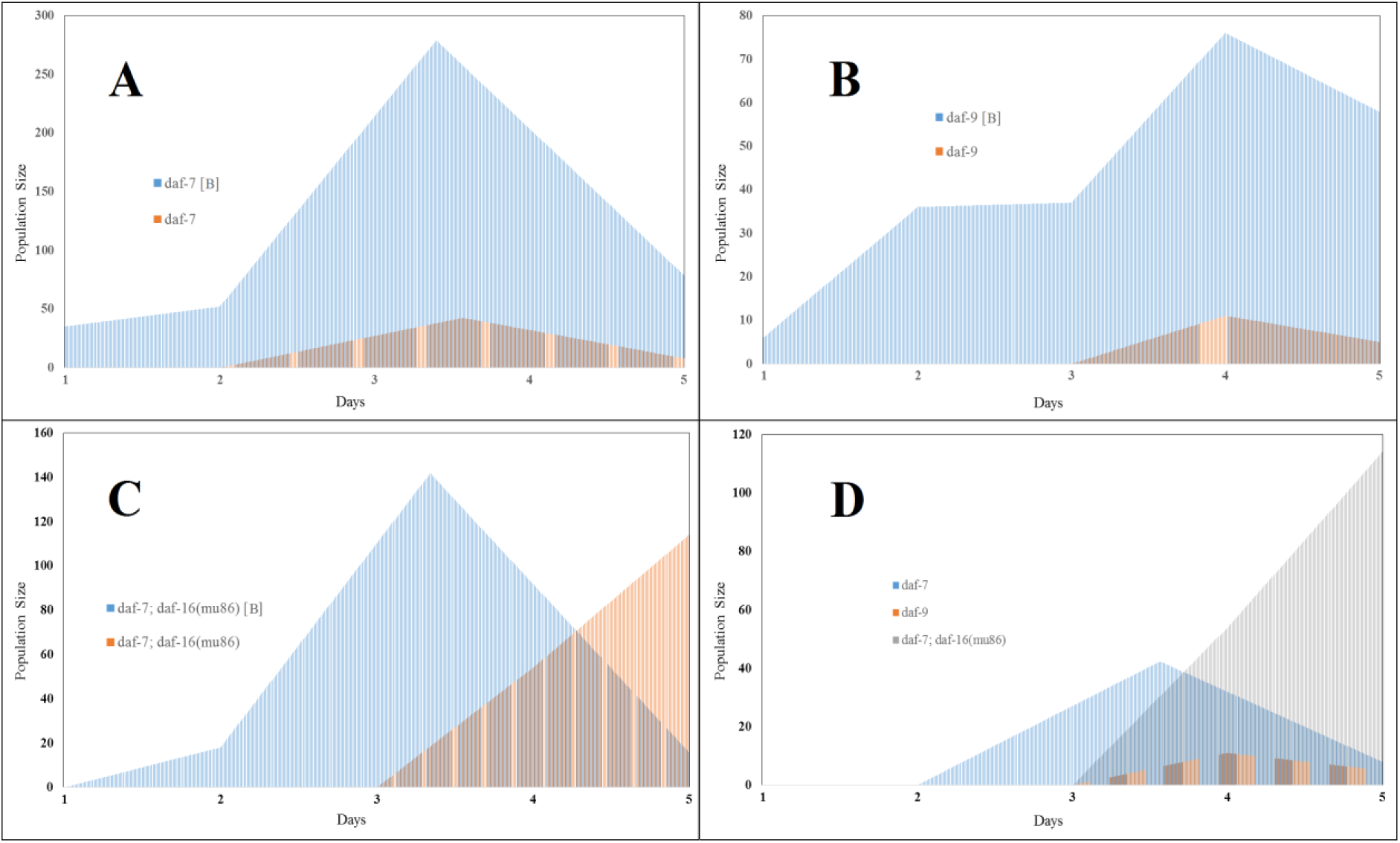
Graphical comparison of the *daf-d* and *daf-c*/*daf-d* genotypes for two environmenta conditions: normal development (blue) versus developmental arrest (orange/pink). A: comparison for daf-7, B: comparison for daf-9, C: comparison for *daf-7*; *daf-16*(mu86), D: comparison of developmental arrest effects for the two *daf-c* and single daf-c/daf-d mutant strains.

**Figure 5.**
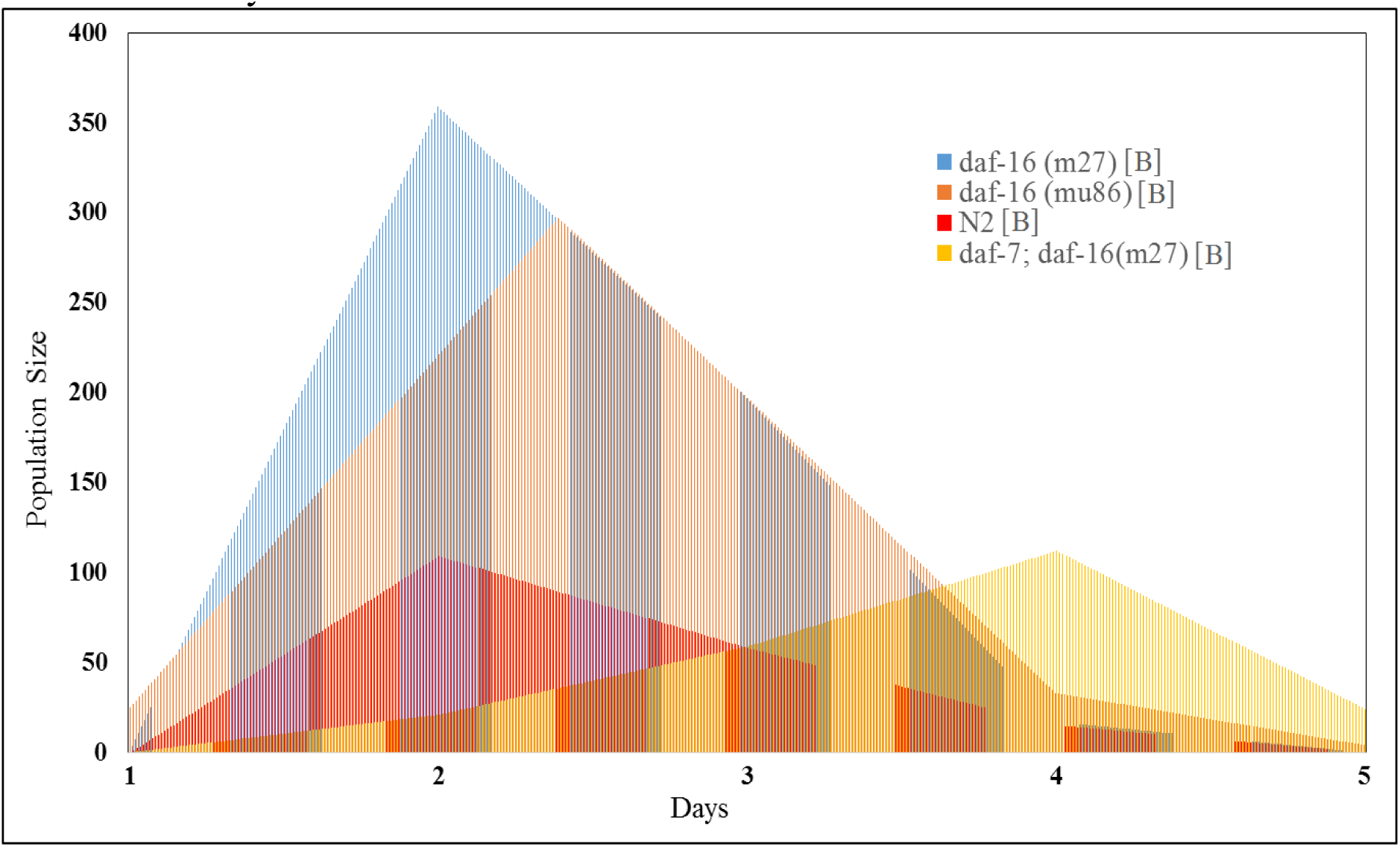
Graphical comparison of baseline reproductive capacity (no extended L1 arrest) for two *daf-d* strains (*daf-16(m27) and daf-16(m27)*), wildtype (N2), and a *daf-c*/*daf-d* strain (*daf-7;daf-16(m27)*).

In contrast to Figure 4, Figure 5 shows the relationships between baseline conditions for wildtype (N2), two *daf-d* mutant strains (*daf-16(m27)* and *daf-16(mu86)*) and a single *daf-c*/*daf-d* mutant (*daf-7*; *daf-16(m27)*). As shown in Table 3, no survivors of extended L1 arrest were observed for either *daf-d* strains nor the *daf-c*; *daf-d* strain represented in Figure 5. Comparing all of the baseline measurements with the baseline for wildtype (N2) allows us to graphically evaluate potential reasons for this failure. In this case, the kurtosis of the interpolated measurement for both *daf-d* (*daf-16*) mutant strains is much higher than wildtype (N2).

The time delay is also either non-existent or very slight, which suggests that mutations of daf-16 at two different alleles enhances reproductive capacity under baseline conditions, but results in very low to nonexistent reproductive robustness. By contrast, the *daf-c*/*daf-d* mutant exhibits little to no change in the kurtosis of peak fecundity relative to the N2 baseline. However, there is a significant delay in peak fecundity, even more pronounced than the delay observed in Figure 4 for reproductive capacity in *daf-c*/*daf-d* genotype after extended L1 arrest. Whether this is connected to reproductive robustness is not clear.

### Developmental Delay Analysis

To better quantify developmental delays resulting from direct developmental perturbation, we can compare the area under the curve (AUC) for different strains and different periods of adult life history. Table S5 shows the AUC for progressive time intervals (e.g. 1-2d, 2-3d) within a single genetic strain/treatment combination. Table S5 also shows the percent difference between the treatment and baseline functions over a specific interval.

There are several examples of intervals over which the treatment condition exceeds the baseline. The generalized lag in development and shift seen in the treatment condition N2 curve (Figure 1) is recognizable in the form of percentages greater than 100% late in the time course (e.g. 4-5d). The large increase in treatment condition *aak-2* is also observed (2230%), but AUC quantification of the *aak-1*; *aak-2* treatment condition curve also reveals that fecundity is a bit higher in both the earliest and latest intervals (109% and 242%, respectively). Recall that these results are not true for the AMPK/daf-c versions of these mutant genotypes. Finally, the *daf-c*/*daf-d* strain *daf-7*; *daf-16(mu86)* also shows a late burst of fecundity which translates treatment condition fecundity above the baseline.

### Starvation Experience over Multiple Generations

While developmental arrest can provide insights into reproductive fitness, investigating this effect experienced over multiple generations can uncover direct evidence of reproductive capacity and robustness with regard to heritability between genetic strains. To make a more explicit connection between development and evolution, the outcomes of starvation experience is assessed in two ways. The first involves potential epigenetic inheritance of information regarding scarce environmental conditions. This by assessing different genetic strains immediately after experiencing starvation (direct) and two generations afterward (indirect). The second way is an assessment of effects due to a combination of repeated starvation and positive selection for fecundity.

### Direct and Indirect Starvation Experience

To understand the immediate and delayed effects of starvation, we can compare pre-starved, direct experience of 5d starvation in the L1 stage, and indirect (2 generations removed) experience of 5d starvation in the L1 stage. Single worms are isolated and allowed to reproduce for 4d, after which the population is counted in bulk. Assessing the population size and effects of experimental treatment in this way provides a way to capture the potential effects of both reproductive efficiency and developmental timing. This survey reproductive capacity is done for both wildtype and various mutant genotypes.

Figure 6 shows the fecundity of single N2 individuals for pre-starvation, starvation-experience, and post-starvation experience conditions. Pre-starvation worms are worms that have no experience with extended L1 arrest. Each measured population originated from a single worm, and the resulting population sizes (after 4d) are represented by blue points. There are two trends associated with the starvation-experience: a smaller population size, and less variation across replicates. In the case of a N2 background, the effect is asymptotic: the decrease in mean population size is quite sharp with starvation-experience, while the decrease after 2 generations is less pronounced.

**Figure 6.**
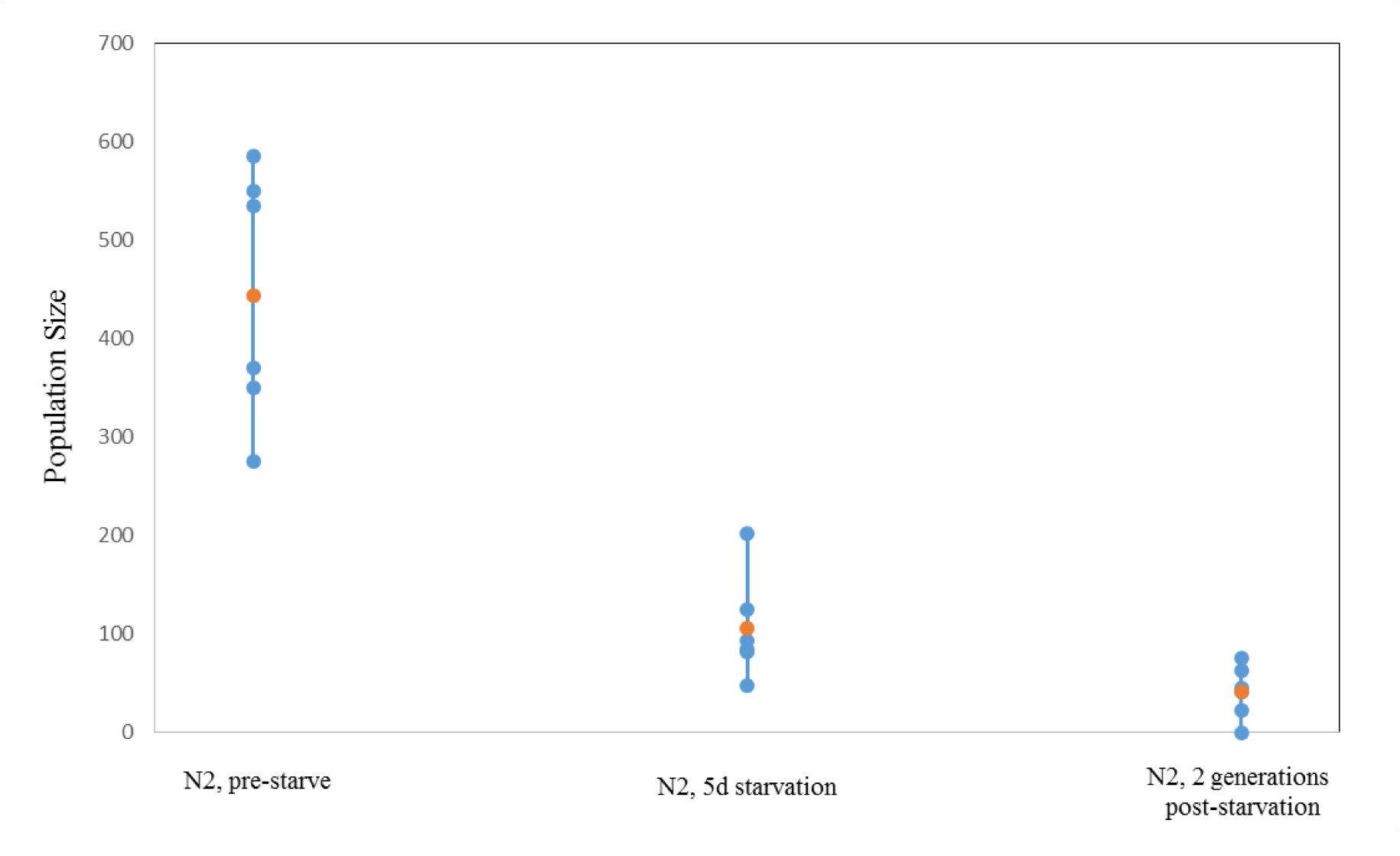
Fecundity (population size) measurements for single wild-type (N2) worms selected from pre-starvation populations (left), individuals that survived 5d extended L1 arrest-related starvation (middle), and individuals two generations removed from those that experienced starvation. Individual populations colored in blue, sample mean colored in orange. All fecundity measurements were taken 4d after each plate was established.

In Figure 7, we can see the immediate and delayed effects of starvation on a series of AMPK mutants: aak-1, aak-2, and aak-1; aak-2. In the case of aak-1 mutants, we observe a less-pronounced effect for starvation-experience as compared to N2, but as is the case with N2 a much smaller population size and reduced variance can be associated with the 2 generations post-starvation experience condition. Figure 7 also shows the immediate and delayed effects of starvation for both the *aak-2* and the *aak-1*; *aak-2* mutants.

**Figure 7.**
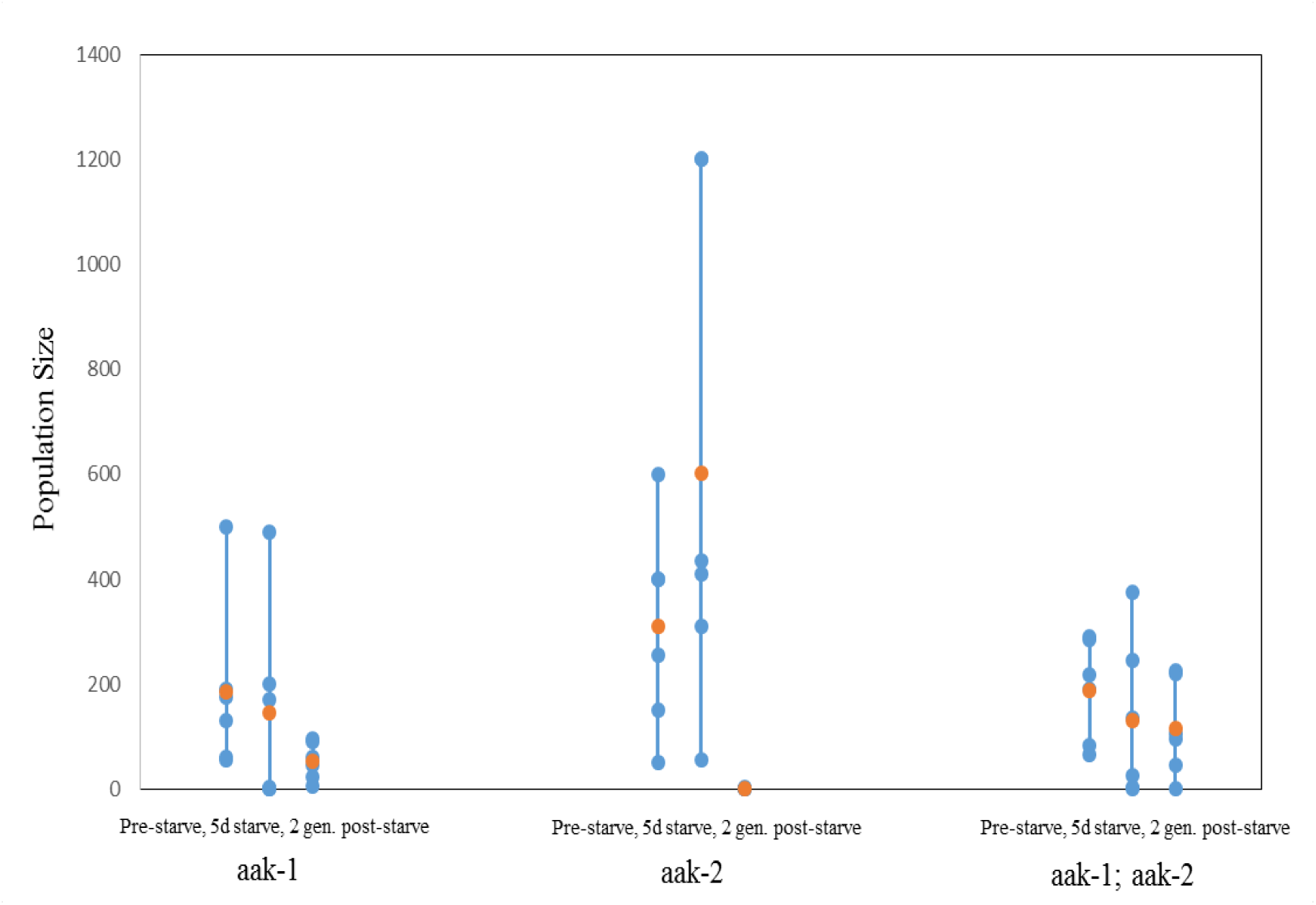
Fecundity (population size) measurements for single AMPK mutant worms selected from pre-starvation populations (left), individuals that survived 5d extended L1 arrest-related starvation (middle), and individuals two generations removed from those that experienced starvation. Individual populations colored in blue, sample mean colored in orange. All fecundity measurements were taken 4d after each plate was established.

In the case of the starvation-experience condition for the single mutant (aak-2), we see greater variation with a tendency towards larger to much larger population sizes. A greater amount of variation between populations is also observed. Contrast this with the post-starvation experience condition, where the populations exhibit a very little variance and small population sizes approaching zero. In the case of the double mutant (*aak-1*; *aak-2*), the mean population size trends progressively downward for both the starvation-experience and post-starvation experience conditions in a manner similar to N2 and *aak-1* mutants. However, the variance between populations increases immediately after starvation-experience and decreases for the poststarvation experience condition.

Figure 8 shows the same set of conditions amongst three AMPK/daf-7 mutant strains: two double mutants (*aak-1*; *daf-7* and *aak-2*; *daf-7*) and a triple mutant (*aak-1*; *aak-2*; *daf-7*). The profile of the *aak-1*; *daf-7* mutant (Figure 3) strain exhibits a decrease across the series of experiences that resembles the N2 genotype. In this case, there is a steep decrease in population size due to starvation, along with a correspondingly large decrease in variance. In comparing starvation-experiencing and post-starvation experiencing individuals, there is a further but slight decrease in both population size and variance between populations.

**Figure 8.**
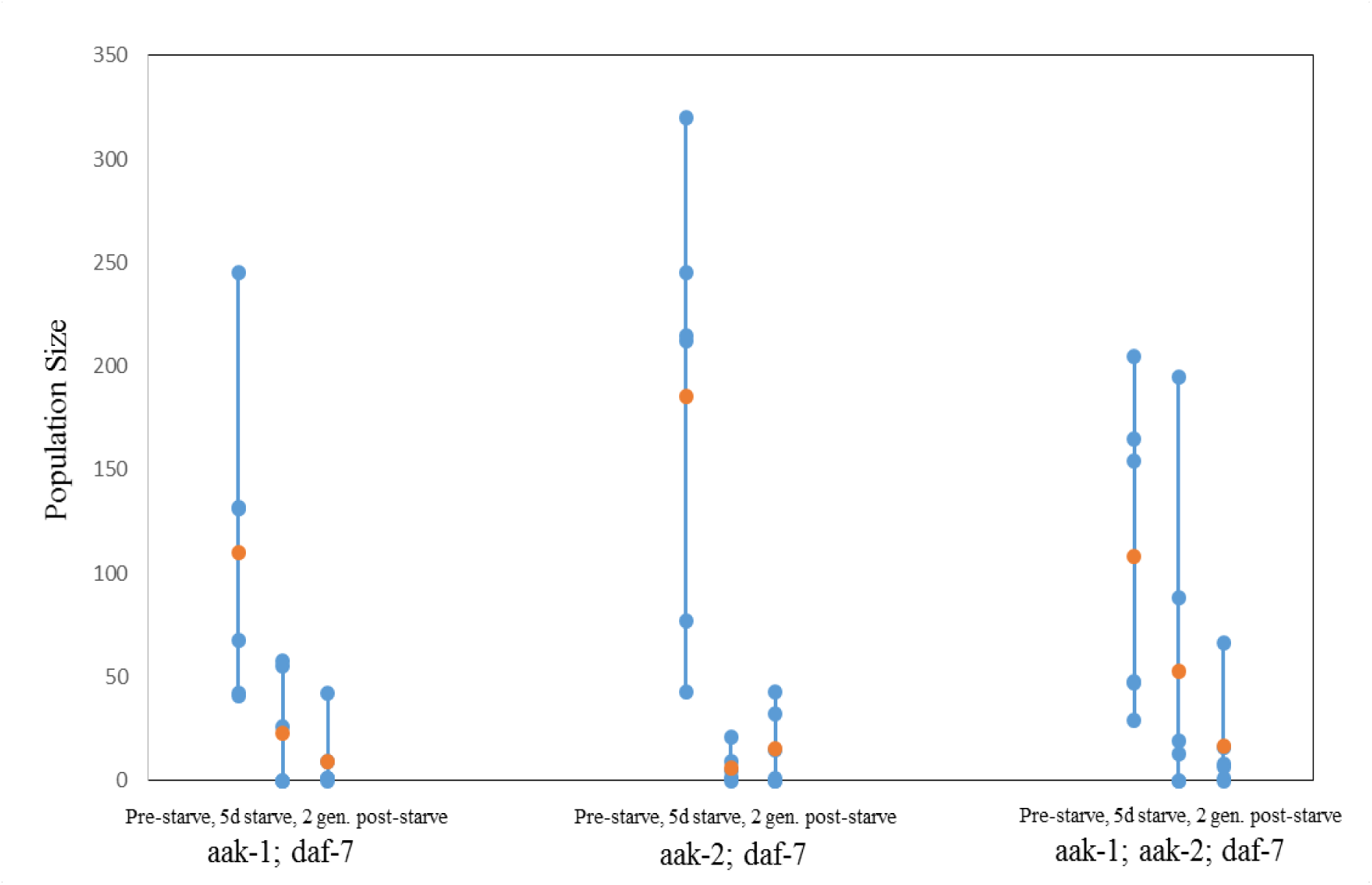
Fecundity (population size) measurements for single AMPK/*daf-c* mutant worms selected from pre-starvation populations (left), individuals that survived 5d extended L1 arrest-related starvation (middle), and individuals two generations removed from those that experienced starvation. Individual populations colored in blue, sample mean colored in orange. All fecundity measurements were taken 4d after each plate was established.

For the triple mutants (*aak-1*; *aak-2*; *daf-7*), we observe a slight but steady decrease in mean population size from pre-starvation to starvation-experiencing to post-starvation experiencing individuals (Figure 8). However, we only see an increase in variance in the starvation-experiencing condition, with a similar amount of variance for the pre-starvation and post-starvation experiencing conditions. By contrast, the *aak-2*; *daf-7* mutants (Figure 8) exhibits a large amount of variation amongst pre-starvation populations. Subsequently (in the starvation-experience condition), there is a large decrease in both mean population size and inter-population variance. In the post-starvation experiencing condition, we observe an increase for both mean population size and variance between populations.

### Starvation Experience Over Evolutionary Epochs

To investigate the longer-term effects of starvation on various mutant genotypes, we conducted a short-term evolution experiment that involves introducing a 5-day starvation period every five generations. The maximum duration of the experiment is three sets of starvation and recovery (16 generations total), although only a few strains survive this long. Individuals generated during the inter-starvation period (the five generations between starvation treatments) are selected twice with respect to reproductive fitness (fecundity).

Figure 9 demonstrates the total number of offspring produced at the second day of reproduction (so-called early reproduction) in the generation immediately following starvation for the three AMPK and three AMPK/daf-7 mutant shown in Figures 2 and 3. An evaluation of early reproduction helps us better understand if there are developmental mechanisms triggered by the repeated exposure to starvation. The producers of offspring immediately following starvation correspond to the first, sixth, and eleventh generation after the beginning of the experiment. We observe uniformity in the number of early offspring amongst *aak-1*, *aak-2*, and *aak-1*; *aak-2* for generations 1 and 6. We also observe a similar uniformity in the number of early offspring amongst *aak-1*; *daf-7*, *aak-2*; *daf-7*, and *aak-1*; *aak-2*; *daf-7* for generations 1.

**Figure 9.**
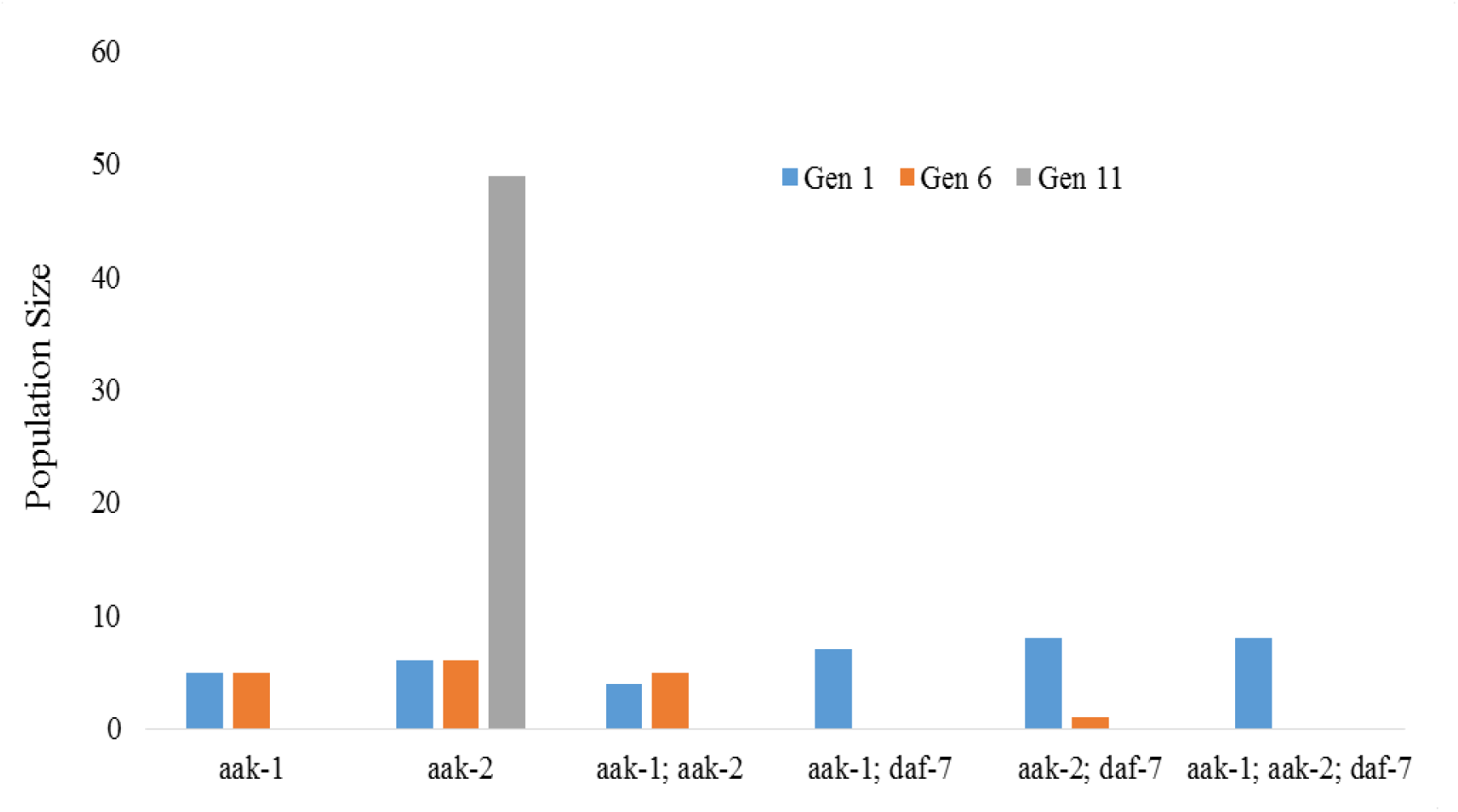
Fecundity (measured as the sum of population size for individual populations) at 2d post-starvation for all AMPK and AMPK/*daf-c* mutants, measured at three different generations(1, 6, and 11).

Generation 11 is where the results exhibit more variability. For the eleventh generation of *aak-2*, there is a large increase in the number of early offspring relative to generations 1 and 6. By contrast, the eleventh generations of *aak-1* and *aak-1*; *aak-2* exhibited no early offspring. Amongst the AMPK-*daf-7* mutants, none of the AMPK; daf-7 mutant strains yielded any survivors. Figure 10 demonstrates how each strain in Figure 4 responds reproductively to starvation on a time-scale of 6d (so-called late reproduction). The population size in Figure 10 includes the number of offspring reported in the 2d measurement. Taken together, Figure 10 provides a sense of the scale of reproduction over that period enabled from a single worm that experienced starvation.

**Figure 10.**
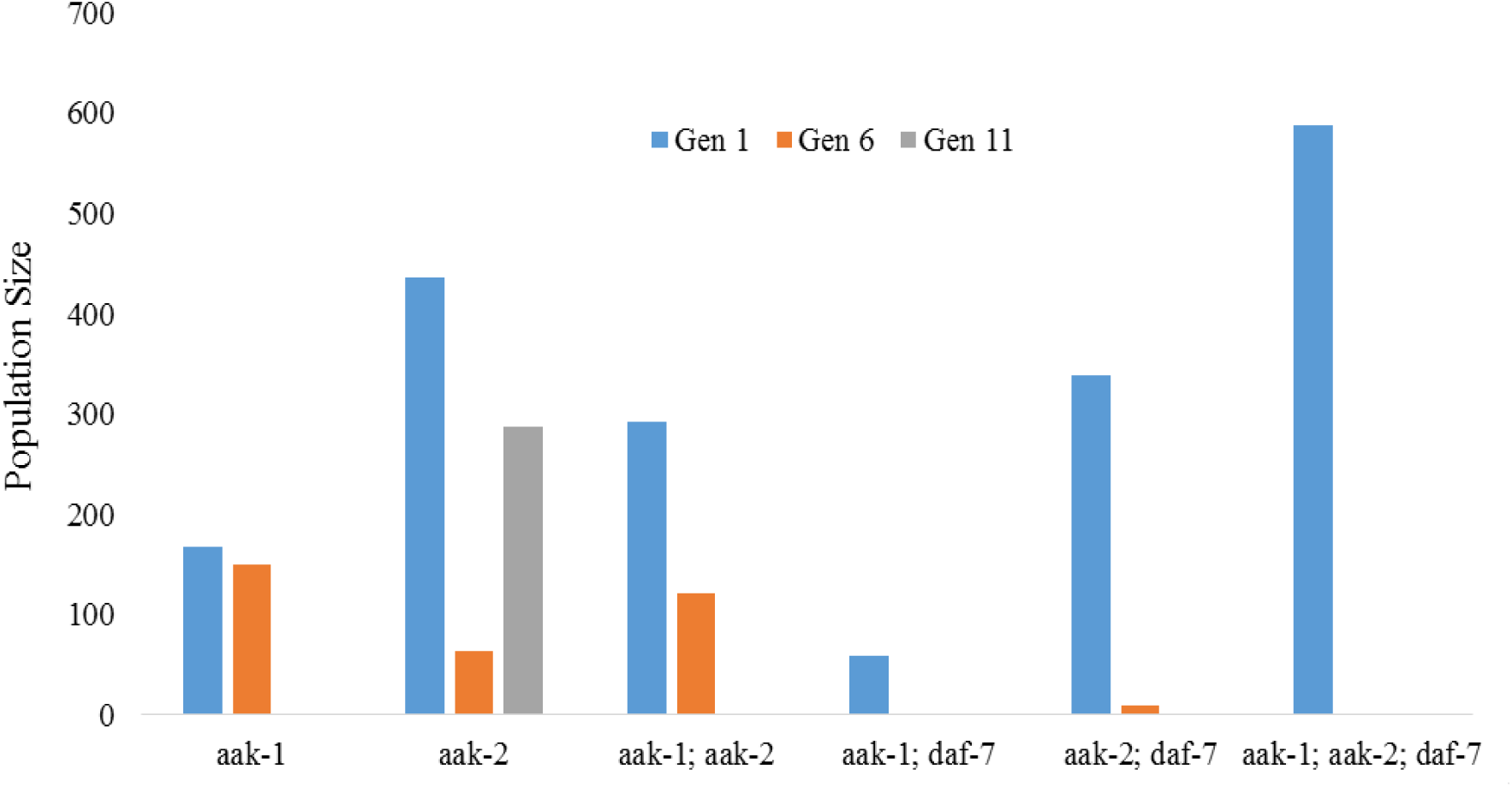
Fecundity (measured as the sum of population size for individual populations) at 6d post-starvation for all AMPK and AMPK/*daf-c* mutants, measured at three different generations(1, 6, and 11).

When the reproductive timescale is extended to 6 days, there is much more variability in population size during generation 1. In particular, both types of *aak-2* and *aak-1*; *aak-2* mutant exhibit much greater population growth (inferred from relative population sizes) than *aak-1*. By generation 6, this variability disappears, and only the 6d results for *aak-1* and *aak-1*; *aak-2* show a sizable increase from the 2d measurements. At generation 11, the only strain to yield survivors is *aak-2*. Whether this represents a long-term compensatory effect is difficult to say, except that the aak-2 was the only AMPK mutant strain to exhibit recovery from starvation at generation 11.

## DISCUSSION

One finding deserving of more explanation is that upon starvation associated with extended L1 arrest, certain AMPK and *daf-c* mutant genotypes demonstrate a compensatory effect. This becomes manifest in the form of enhanced reproductive capacity, which may occur as a result of shifts in developmental timing and other genomic mechanisms related to heterochronic progression. We may be observing a set of compensatory mechanisms in the *aak-2* and *aak-2*; *daf-7* mutants triggered by the experience of starvation. There may also be a compensatory effect in the AMPK double mutant strains (aak-1; aak-2 and aak-1; aak-2; daf-7) as well. Mutants of the AMPK pathway is involved in both the aging and metabolism functions of C. elegans. In particular, a gene expression analysis by Shin et.al (2011) shows that aak-2 mutants exposed to stress exhibit a larger number of upregulated and a smaller number of downregulated genes relative to unstressed aak-2 mutants. In general, there is a larger number of differentially-regulated genes between aak-2 mutants and wildtype (N2), providing some evidence for a compensatory mechanism.

Another finding demonstrated by the extended L1 arrest and multigenerational data is that *daf-c* mutant genotypes show rather pronounced reproductive delay, while *daf-d* mutant genotypes show no ability to survive the extended L1 arrest treatment. Due to the combination of techniques, this is shown both within and between generations. While this is less pronounced in other types of mutant, and even the wildtype strain, perturbation of developmental timing and selection for fecundity seem to have convergent effects.

In terms of the multigenerational experiments, there are a number of caveats in the presented results. Perhaps the biggest involves the reproductive nature of *C. elegans* (extensive and repeated hermaphroditic backcrossing) can impact the reproductive robustness of a population put under selection. Namely, the impact of selection for fecundity can lead to linkage disequilibrium, which in some cases can lead to the linkage of deleterious and advantageous alleles (Thomas, Woodruff, and Haag, 2012). In the results of the natural selection experiments shown here, populations undergo both selection and expansion. This not only minimizes the deleterious effects of selection, but also allows us to select for epigenetic mechanisms, which result in positive fecundity in starvation-experiencing populations.

When reproductive capacity is high in the face of experiencing extended L1 arrest, the genotype in question is robust to developmental stress. However, in cases where reproductive capacity is low, the genotype in question is fragile to developmental stress. This is a product of both individuality and how the individual’s genotype buffers them from the starvation experience. It is of note that the extended arrest period produces two outcomes: variation in the number of survivors per treatment, and variation in their subsequent developmental trajectory. There is a necessarily large amount of variation with respect to the number of eggs used to create a population of synchronized L1 worms. As the mortality rate over the extended arrest period differs from strain to strain, and the sampling procedure to create isolates is random, we could be selecting either super-robust individuals or individuals with no special ability to survive and recover from starvation conditions. Even though the genomic mechanism behind this individuality is unclear, it may play a role in within-strain variation, particularly the putative compensatory mechanisms described earlier.

The same holds true for variation in the intra-strain rate of recovery. Anecdotal observations suggests that the time from late L1 to the L4 stage can range from 24h post return to food to >48h. Amongst the daf-c, daf-c/daf-d crosses, and AMPK, daf-c crosses, there were several cases of delayed adulthood. A few individuals also exhibited an extended larval period without surviving to reproductive age. While this study focuses on the reproductive period post-recovery, particularly on randomly sampling the survivors of the process, methods do exist for calculating survival rates for populations undergoing extended L1 arrest (Artyukhin, Schroeder, and Avery, 2013; Jobson et.al, 2015).

Yet how can we use the concept of developmental robustness to characterize the flexibility of developmental pathways in specific genotypic strains? Wagner (1996) and Ciliberti, Martin, and Wagner (2007) use complex networks of epigenetic interactions and gene co-expression data, respectively to build an analyzable model of flexibility due to adaptation. In both cases, we find that networks for similar phenotypic processes can lead to many different patterns of connectivity in the network. This set of alternative pathways is analogous to decanalization, and points to changes that do not require adaptive mutations. These neutral changes in network topology represents the degree of robustness in the system. For example, the number of equivalent patterns of connectivity increases, the more robust to developmental disruptions (e.g. starvation) the system becomes (Ciliberti, Martin, and Wagner, 2007). In the case of comparing the robustness of specific mutant genotypes to starvation, we should expect some mutations to render dysfunctional gene expression patterns, resulting in a slower or unpredictable expansion of the neutral set of possibilities.

One way we can infer the way these spaces might vary with respect to genotype is by looking at the KDE distributions in Figure S1. In some classes of mutant, there are very clear shifts in fecundity relative to the wildtype. These shifts might reflect the underlying adaptive capacity driven by functional loss or functional compensations of the underlying gene networks. Another way we can infer the expansion of these spaces is by identifying unexpected patterns in the data. While our sample size is small, there are nevertheless genotypic-specific results that stand out from more general trends exhibited by both mutant and wildtype genotypes. Future work would involve confirming these interpretations using empirical and/or theoretical interaction networks for various functional mutations for varying degrees of perturbation.

## ACKNOWLEDGEMENTS

We would like to thank Dr. Nathan Schroeder, Rebecca Androwski, Kristen Flatt for their advice on *C. elegans* classical genetics techniques and worm culture. I would also like to thank viewers of poster 766C at the 20^th^ International *C. elegans* Meeting for their comments. Some of the work presented in this paper was supported by the National Institute of General Medical Sciences of the National Institutes of Health (Award Number IR01GM111566-01). Some strains were provided by the CGC, which is funded by NIH Office of Research Infrastructure Programs (P40 OD010440).

## METHODS

All data (baseline and extended L1 arrest, reproductive time-series and transgenerational population counts) can be found on Figshare at doi:10.6084/m9.figshare.3122113.

Organisms:

### Strains

Eight (8) strains were acquired from either the Caenorhabditis Genomics Center (CGC, http://cbs.umn.edu/cgc/) or Nathan Schroeder’s Laboratory at University of Illinois Urbana-Champaign. These genotypes were: *qIs56[Plag-2::GFP]; daf-7(e1372), qIs56[Plag-2::GFP]; daf-9(rh50), N2, myIs14[Pklp-6::GFP]; daf-7(e1372); daf-16(mu86), daf-16(m27), qIs56[Plag-2::GFP], lag-2(q420),* and *daf-16(mu86)*. Further information regarding the [Plag-2::GFP] and [Pklp-6::GFP] constructs can be found in Schroeder et.al (2013).

### Genetic Crosses

Seven (7) additional crosses were constructed to gain additional data about the contributions of the AMPK and *daf-7* mutant genotypes to evolvability and developmental plasticity. These genotypes were: *qIs56[Plag-2::GFP]; daf-7(e1372); daf-16(m27), qIs56[Plag-2::GFP]; aak-1(tm1944); daf-7(e1372), qIs56[Plag-2::GFP]; aak-2(ok524); daf-7(e1372), qIs56[Plag-2::GFP]; aak-1(tm1944); aak-2(ok524); daf-7(e1372), qIs56[Plag-2::GFP]; aak-1(tm1944), qIs56[Plag-2::GFP]; aak-2(ok524), qIs56[Plag-2::GFP]; aak-1(tm1944); aak-2(ok524)*. More details on the design can be found on Figshare at doi: 10.6084/m9.figshare.2436289.v1

Analyses and Techniques:

### Graphs

All graphs and data presentation were made in Excel, MATLAB, and R. All statistical analyses were run in MATLAB and R.

### Growth Conditions

All strains maintained at 25C except for the *lag-2* mutant (15C), and *daf-7* and *daf-9* mutants (22C).

### Sample Size

For both the baseline and extended L1 arrest conditions, seven (7) individuals were isolated for each strain and measured over the course of five (5) days. This provided *N* = 7 and *n* = 35 for each strain*condition combination. For the transgenerational data, eight (8) individuals were isolated per strain. For the secondary isolated lines (see transgenerational selection), the number of selected populations is contingent upon the distribution of population sizes at 6d.

### Transgenerational Selection

For the transgenerational data, each strain is starved for 5d and then plated. After 24-48h, eight (8) L4 individuals were isolated per strain. In cases where eight individuals could not be recovered, the maximum number of individuals were isolated. At 6d, anywhere from 1-4 of the isolated lines with the largest population sizes were selected for continuation, by isolating three (3) individuals from each of the selected isolated lines. Each of the selected individuals were used to found a secondary isolated line. The offspring of these selected lines are harvested and subject to another round of extended L1 arrest. This process is replicated a maximum of three times, although not all strains survived the first and second round. The secondary isolated lines are counted after 5d, and the 1-2 most fecund sets of secondary isolated lines were harvested, pooled, and re-exposed to the extended L1 arrest protocol. Where possible, the procedure is repeated to obtain data for the 6th and 11th generation.

### Extended L1 Arrest

Worms from single plates that contain gravid worms are suspended in M9 buffer and transferred to an Eppendorf tube. Worms are centrifuged for 8 minutes at 4000g. The supernatant is removed and pellet is suspended in a Bleach/NaOH solution (55% dH2O, 25% NaOH at 1% concentration, and 20% Bleach). Mixture is mixed by inversion and rotation for 10 minutes. Eppendorf tubes are then centrifuged again for 8 minutes at 4000g. The supernatant is removed, and pellet is rinsed 2x in buffer: the first time in M9 buffer with 0.01% Triton, and the second time in M9 buffer alone. He remaining pellet (consisting of unhatched eggs) is suspended in 4mL M9 buffer. The tube is then stored overnight (to hatch the eggs into L1 stage) on an incubator shaker at 50RPM and a strain-appropriate temperature. Tubes are then stored for 5d at 15C. After 5d, the tube is centrifuged for 10 minutes at 4000g. The resulting pellet is then suspended in 500uL of M9 buffer are transferred to a plate filled with NGM agar is seeded with 100uL of OP50 media. At 24-48h post transfer, plates are then surveyed for survivors. Individuals are then picked and transferred to new plates for measurement.

### Determining Population Size

For every generation, five replicates were created. After a 60mm diameter plate filled with NGM agar is seeded with 100uL of 0P50 media and allowed to dry, a single L4 stage worm is placed on each of the five plates. Each plate is allowed to grow for 3d at optimal temperatures for the given strain. At the end of the 3d period, plates are examined for offspring and counted. Anywhere from 1-3 plates are then selected to represent the subsequent generation. Five new plates are seeded from the selected populations with L4 stage representatives in proportion to their population size (e.g. larger populations get more representatives in the next generation).

Selection criterion:

## Hanging Drop method

The hanging drop method was proposed by Muschiol, Schroeder, and Traunspurger (2009) as a means of assessing the fecundity of a single worm over a discrete period of time isolated from maternal effects and with minimal counting error. The hanging drop is conducted by harvesting a series of worms at the L4 stage and plating them one worm per plate. Every 24h, each plate is checked for offspring. The number of offspring are counted, and the parent is transferred to a new plate. This was done over the course of 3d for each evolved strain to serve as a control (baseline). The population count for each strain was derived by summing all three daily measurement per replicate and then averaging across the replicates.

## Kernal Density Estimation (KDE) of Reproductive diversity

The KDE analysis was implemented and graphed in MATLAB. Classes of genotype (see Methods, Organisms for specific alleles) were defined as: wildtype (N2, *qIs56*), NOTCH (*lag-2* mutant), AMPK (*aak-1*, *aak-2*, *aak-1*; *aak-2* mutants), AMPK/*daf-c* (*aak-1; daf-7, aak-2; daf-7, aak-1; aak-2; daf-7* mutants), *daf-c* (*daf-7, daf-9), daf-d (daf-16(m27), daf-16(mu86)* mutants), and *daf-c/daf-d* (*daf-7; daf-16(m27), daf-7; daf-16(mu86)* mutants). Net fecundity is measured by subtracting the per-day growth of non-starved individuals starting at the L4 stage from similar individuals that have experienced extended L1 arrest. Both sets of growth measurements were taken using the Hanging Drop method. The frequency of the resulting fecundity measurements were then plotted on a histogram.

## Genotyping for Mutant Construction

Genotyping for the aak-1(tm1944) and aak-2(ok524) mutants were done using the following primers:tm1944; internal (TCACACGTCTCTTCCGTGTT), left flanking (TCGCGTCCAGAAGAAGATTT), right flanking (TCCCTTTCTTCGCTCACTTT). ok524; internal (CAAAGTCCGCAATCTTCACA), left flanking (TCATCCGCCTCTACCAAGTC), right flanking (TCAAATCCCATTTCGCTTTC). Sequences for primer design retrieved from Genbank http://www.ncbi.nlm.nih.gov/. Primer design conducted using Primer3 (http://biotools.umassmed.edu/primer3/primer3web_input.htm). Using Blastn, primers were evaluated (e.g. E-value) for similarity with bacterial sequences and other Nematode sequences.

## Quantitative Measures

### Interpolation of Fecundity Measurements

The fecundity measurements collected via the Hanging Drop method were resampled using a linear interpolation method. The objective was to transform a sparsely sampled time series into a continuous curve for statistical comparisons. These curves were calculated for both the baseline and treatment conditions. The curves also produced a characteristic peak when plotted in a bivariate space. The linear interpolation method can be expressed as

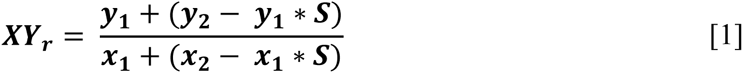

where *XY*_*r*_ is the resampled function over interval *r*, *x*_*n*_ is a specific time point *n*, *y*_*n*_ is the measured fecundity at time point *n*, *S* is the sampling interval (0.01 days).

### Fold-Change Measurement

To compare two linearly interpolated curves in a pairwise manner, the fold-change in both the time of peak fecundity (maximum on the x-axis) and number of individuals representing peak fecundity (maximum on the y axis) was calculated. The fold-change for a pairwise comparison can be calculated as

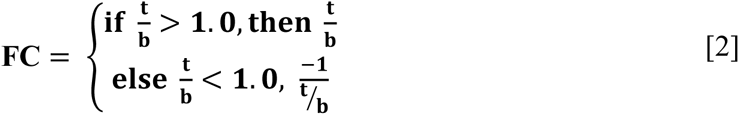

### Area Under the Curve (AUC) Intervals

AUC is calculated from the linearly interpolated fecundity data using the trapezoidal method. The sampling interval was 0.01 time units. The trapezoidal method can be defined as

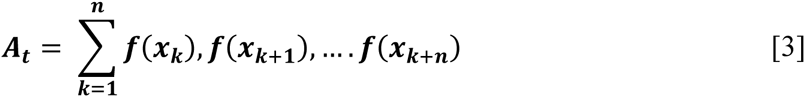

where *A*_t_ is the AUC for a specific time interval, *x*_*k*_ is the AUC for a single temporal sampling point, and *n* is the size of the sampling interval. For a single day, *n* = 100.

## REFERENCES

Apfeld, J., O’Connor, G., McDonagh, T., DiStefano, P.S, and Curtis, R. 2004. The AMP-activated protein kinase AAK-2 links energy levels and insulin-like signals to lifespan in *C. elegans*. Genes and Development, 18, 3004-3009.

Artyukhin, A.B. and Avery, L. 2013. L1 arrest affects dauer decisions in *C. elegans*. Worm Breeder’s Gazette, 19(4), 19-20.

Artyukhin, A.B., Schroeder, F.C., and Avery, L. 2013. Density dependence in *Caenorhabditis* larval starvation. Scientific Reports, 3, 2777.

Baugh, L.R. 2013. To grow or not to grow: nutritional control of development during *Caenorhabditis elegans* L1 arrest. Genetics, 194(3), 539-555.

Burnell, A.M., Houthoofd, K., O’Hanlon, K., and Vanfleteren, J.R. 2005. Alternate metabolism during the dauer stage of the nematode *Caenorhabditis elegans*. Experimental Gerontology, 11, 850-856.

Ciliberti, S., Martin, O.C., Wagner, A. 2007. Robustness Can Evolve Gradually in Complex Regulatory Gene Networks with Varying Topology. PLoS Computational Biology, 3(2), e15.

Fukuyama, M., Sakuma, K., Park, R., Kasuga, H., Nagaya, R., Atsumi, Y., Shimomura, Y., Takahashi, S., Kajiho, H., Rougvie, A., Kontani, K., and Katada, T. 2012. *C. elegans* AMPKs promote survival and arrest germline development during nutrient stress. Biology Open, 1, 929-936.

Harvey, S.C. and Orbidans, H.E. 2011. All Eggs Are Not Equal: the maternal environment affects progeny reproduction and developmental fate in *Caenorhabditis elegans*. PLoS One, 6(10), e25840.

Jobson, M.A., Jordan, J.M., Sandrof, M.A., Hibshman, J.D., Lennox, A.L., and Baugh, L.R. 2015. Transgenerational Effects of Early Life Starvation on Growth, Reproduction and Stress Resistance in *Caenorhabditis elegans*. Genetics, 201(1), 201-212.

Johnson, T.E., Mitchell, D.H., Kline, S., Kemal, R., and Foy, J. 1984. Arresting development arrests aging in the nematode *Caenorhabditis elegans*. Mechanisms of Ageing and Development, 28, 23–40.

Kang, C., and Avery, L. 2009. Systemic regulation of starvation response in *Caenorhabditis elegans*. Genes and Development, 23, 12–17.

Lee, I., Hendrix, A., Kim, J., Yoshimoto, J., You, Y-J. 2012. Metabolic Rate Regulates L1 Longevity in *C. elegans*. PLoS One 7(9), e44720.

Maxwell, C.S., Antoshechkin, I., Kurhanewicz, N., Belsky, J.A., and Baugh, L.R. 2012. Nutritional control of mRNA isoform expression during developmental arrest and recovery in *C. elegans*. Genome Research, 22, 1920–1929.

Michaelson, D., Korta, D.Z., Capua, Y., and Hubbard, E.J. 2010. Insulin signaling promotes germline proliferation in *C. elegans*. Development, 137(4), 671-680.

Moczek, A.P., Sultan, S., Foster, S., Ledon-Rettig, C., Dworkin, I., Nijhout, H.F., Abouheif, E., and Pfennig, D.W. 2011. The role of developmental plasticity in evolutionary innovation. Proceedings of the Royal Society B, 278, 2705-2713.

Mukhopadhyay, A. and Tissenbaum, H.A. 2007. Reproduction and longevity: secrets revealed by *C. elegans*. Trends in Cell Biology, 17(2), 65-71.

Murphy C.T., Hu P.J. 2013. Insulin/insulin-like growth factor signaling in *C. elegans*, WormBook (December 26). In “WormBook”, The *C. elegans* Research Community eds., WormBook, doi/10.1895/wormbook.1.164.1, http://www.wormbook.org.

Muschiol, D., Schroeder, F., and Traunspurger, W. (2009). Life cycle and population growth rate of *Caenorhabditis elegans* studied by a new method. BMC Ecology, 9, 14.

Rechavi, O., Houri-Ze’evi, L., Anava, S., Siong Sho Goh, W., Kerk, S.Y., Hannon, G.J., Hobert, O. 2014. Starvation-induced transgenerational inheritance of small RNAs in *C. elegans*. Cell 158: 277-287.

Schroeder, N.E., Androwski, R.J., Rashid, A., Lee, H., Lee, J., Barr, M.M. 2013. Dauer-specific dendrite arborization in *C. elegans* is regulated by KPC-1/Furin. Current Biology, 23(16), 1527-1535.

Shin, H., Lee, H., Fejes, A.P., Baillie, D.L., Koo, H-S., and Jones, S.J.M. 2011 Gene expression profiling of oxidative stress response of *C. elegans* aging defective AMPK mutants using massively parallel transcriptome sequencing. BMC Research Notes, 4, 34.

Thomas, C.G., Woodruff, G.C., and Haag, E.S. 2012. Causes and consequences of the evolution of reproductive mode in *Caenorhabditis nematodes*f. Trends in Genetics, 28(5), 213-220.

Vowels J.J. and Thomas, J.H. 1992. Genetic Analysis of Chemosensory Control of Dauer Formation in *Caenorhabditis elegans*. Genetics, 130, 105-123.

Wagner, A. 1996. Does Evolutionary Plasticity Evolve? Evolution, 50(3), 1008-1023.

West-Eberhard, M.J. 2003. Developmental Plasticity and Evolution. Oxford University Press, Oxford, UK.

Wolkow C.A., Kimura K.D., Lee M.S., and Ruvkun G. 2000. Regulation of *C. elegans* life-span by insulin-like signaling in the nervous system. Science, 290, 147–150.

Zaslaver, A., Baugh, L.R., and Sternberg P.W. 2011. Metazoan operons accelerate recovery from growth-arrested states. Cell, 145, 981–992.

